# Dynamic methylome modification is associated with mutational signatures in aging and the etiology of disease

**DOI:** 10.1101/2020.12.22.423778

**Authors:** Gayatri Kumar, Kashyap Krishnasamy, Naseer Pasha, Naveenkumar Nagarajan, Bhavika Mam, Mahima Kishinani, Vedanth Vohra, Renuka Jain, Villoo Morawala Patell

## Abstract

Reversible epigenetic changes within the loci of genes that regulate critical cell processes have recently emerged as important biomarkers of disease pathology. It is then natural to consider the consequences for population health risk of such epigenetic changes during the aging process. Specifically, the interplay between dynamic methylation changes that accompany aging and mutations that accrue in an individual’s genome over time need further investigation. The current study investigated the role of dynamic methylation acting together with gene variants in an individual over time to gain insight into the evolving epigenome–genome interplay that affects biochemical pathways controlling physiological processes during aging.

We completed a whole-genome methylation and variant analysis in a non-smoking Zoroastrian-Parsi individual, collecting two samples, 12 years apart (at 53 and 65 years respectively) (ZPMetG-Hv2a-1A (old, t_0_), ZPMetG-Hv2a-1B (recent, t_0_+12)) and analysing them using a GridION Nanopore sequencer at 13X genome coverage overall. We further identified the single nucleotide variants (SNVs) and indels in known CpG islands by employing the Genome Analysis Tool Kit (GATK) and MuTect2 variant-caller pipeline with the GRCh37 (patch 13) human genome as reference.

We found 5258 disease-relevant genes that had been differentially methylated in this individual over 12 years. Employing the GATK pipeline, we found 24,948 genes, corresponding to 4,58,148 variants, specific to ZPMetG-Hv2a-1B, indicating the presence of variants that had accrued over time. A fraction of the gene variants (242/24948) occurred within the CpG regions that were differentially methylated, with 67/247 exactly coincident with a CpG site. Our analysis yielded a critical cluster of 10 genes that were each significantly methylated and had variants at the CpG site or the ±4 bp CpG region window. Kyoto Encyclopedia of Genes and Genomes (KEGG) enrichment network analysis as well as Reactome and STRING analysis of gene-specific variants indicated an impact on biological processes regulating the immune system, disease networks implicated in cancer and neurodegenerative diseases, and transcriptional control of processes regulating cellular senescence and longevity. Additional analysis of mutational signatures indicated a majority of C>T transitions followed by T>C transitions in the more recent sample, ZPMetG-Hv2a-1B.

Our current study provides additional insight into the aging methylome over time and the interplay between different methylation and gene variants in the etiology of disease.

## Introduction

Aging is a complex and time-dependent deterioration of physiological processes. Increased human life expectancy has resulted in higher morbidity rates, as advanced age is the predominant risk factor for several diseases, including cancer, dementia, diabetes, and cardiovascular disease (CVD)^1^. In addition to molecular and cellular factors, such as cellular senescence and telomere attrition, epigenetic changes that govern physiological processes comprise a significant component of the aging process^2^. Methylation signatures specifically influence gene expression without altering the DNA sequence, connecting intrinsic and extrinsic signals. The most common methylation modifications involve the transfer of a methyl (CH_3_) group from *S*-adenosyl methionine (SAM) to the fifth position of cytosine nucleotides, forming 5-methylcytosine (5mC)^3^. Methylation is a dynamic event, in which reversal occurs at certain sites, while progression with age causes methylation at many CpG sites in intergenic regions, such as transcription start sites (TSSs^4^). The interplay between epigenetic events is manifested in the regulation of major biological processes^5^, such as development, differentiation, genomic imprinting, and X chromosome inactivation (XCI).

DNA methylation, both heritable and dynamic, exhibits a strong correlation with age and age-related outcomes. Additionally, the epigenome responds to a broad range of environmental factors and is sensitive to environmental influences^6^, including exercise, stress^7^, diet^8^, and sleep patterns. Epigenetic profiling of identical twins^9^ indicates that methylation events are independent of genome homology, with global DNA methylation patterns differing between older and younger twins, consistent with an age-dependent progression^10^ of epigenetic changes. Global methylation changes over an 11-year span in participants of an Icelandic cohort, as well as age- and tissue-related alterations in certain CpG islands from an array of 1413 arbitrarily chosen CpG sites near gene promoters, further corroborate the evidence for dynamic methylation patterns over time^11^. Promoter hypermethylation has been shown to increase mutation rates, suggesting the influence of epigenetics on genetics. These studies support the role of epigenome alterations in aging-associated genetic changes. It has been reported that genes related to various tumors can be silenced by heritable epigenetic events involving chromatin remodeling or DNA methylation (DNAm). In addition to dynamic methylation changes over time, recent studies have also suggested that epigenetic markers, and their maintenance, are controlled by genes and are tightly linked with DNA variants^12^ that accrue in an individual genome over time. In fact, the methylation of cytosine to 5mC in the coding regions of genes increases the probability of mutations and consequent spontaneous hydrolytic deamination resulting in C>T transitions.

The epigenetic milieu therefore provides a homeostatic mechanism at the molecular level that allows phenotypic malleability in response to the changing internal and external environments. The flexibility and dynamism of the methylome, as monitored through epigenetic profiling, suggests the importance and influence of lifestyle changes in the control of gene regulation. This also makes methylome studies an attractive means to identify epigenetic hotspots and evaluate the impact of lifestyle changes on aging as well as provide potential targets for therapeutic intervention^13,14^. Besides methylation changes, mutational signatures, including somatic variations, can change the preponderance of methylation of CpG islands. CpG–SNP interactions in the promoter regions have been found to be associated with various disorders, including type 2 diabetes^15^, breast cancer^16^, coronary heart disease^17^, and psychosis^13^.

Therefore, comparison of comprehensive methylation patterns in healthy individuals at multiple time points can determine whether such changes are early indicators of a late-onset chronic disease. Identifying such indicators earlier can improve disease prognosis, with the potential for reversibility. Our current epigenetic study is unique in analysing genome– epigenome interactions from an individual of the dwindling endogamous Zoroastrian Parsi community, which has a higher median life span and, as a result, a higher incidence of aging-associated conditions, including neurodegenerative conditions, such as Parkinson’s disease; cancers of the breast, colon, gastrointestinal tract, and prostrate; auto-immune disorders; and rare diseases.

To identify age- and disease-associated epigenetic changes in the Parsi population, we performed an intra-individual methylome profile of a Zoroastrian-Parsi individual with Nanopore-based sequencing technology using samples collected 12 years apart. The two samples, ZPMetG-HV2a-1A (old) and ZPMetG-HV2a-1B (recent) (12 years apart) were sequenced using Oxford Nanopore Technology (ONT). Comparing the samples, we found that aging increased global methylation frequencies for the equivalent CpG sites, with an overall increase of hypermethylated regions across genic regions compared with non-genic regions. Some genes (108) were significantly hypermethylated, while 304 genes were significantly hypomethylated. We also identified 10 unique CpG–SNP interactions in the form of variants occurring at a CpG site and across the CpG region that may affect the biological processes encoded in the genic regions. The enrichment in certain genes implicated pathways involved in the immune response, cancer, neurogenerative diseases, and physiological processes regulating cellular proliferation, senescence, aging, and longevity.

## Materials and Methods

### DNA Extraction

High-molecular-weight (HMW) genomic DNA was isolated from the buffy coat of EDTA-treated whole blood using the Qiagen Whole Blood Genomic DNA extraction kit (cat. #69504). The Extracted DNA Qubit™ dsDNA BR assay kit (cat. #Q32850) was used in preparation for the Qubit 2.0® Fluorometer (Life Technologies™).

### DNA QC

HMW DNA of optimal quality is a prerequisite for Nanopore library preparation. The quality and quantity of the gDNA was estimated using a Nanodrop spectrophotometer and a Qubit fluorometer using the Qubit™ dsDNA BR assay kit (#Q32850) from Life Technologies, respectively.

### DNA purification

The DNA was subjected to column purification using the Zymoclean Large Fragment DNA Recovery kit (Zymo Research, USA), followed by fragmentation using g-TUBE device (Covaris, Inc.). The purified DNA samples were used for library preparation.

### Library preparation

A total of 2 μg from each sample was used for Nanopore library preparation using the Nanopore Ligation Sequencing kit (cat. #SQK-LSK109, Oxford Nanopore Technology, Oxford, UK). Briefly, 2 μg of gDNA from each sample was end-repaired using an NEBnext Ultra II End Repair kit, (New England Biolabs, MA, USA) and purified using 1x AmPure beads (Beckman Coulter, USA). Adapter ligation (AMX) was performed at RT (20 °C) for 20 min using NEB Quick T4 DNA ligase (New England Biolabs, MA, USA). The reaction mixture was purified using 0.6X AmPure beads (Beckman Coulter, USA), and the sequencing library was eluted in 15 μl of elution buffer provided in the ligation sequencing kit.

### Sequencing and sequence processing

Sequencing was performed on a GridION X5 sequencer (Oxford Nanopore Technologies) using a SpotON flow cell R9.4 (cat. #FLO-MIN106) as per the manufacturer’s recommendation. Nanopore raw reads (“fast5” format) were base called (“fastq5” format) using Guppy v2.3.4 software.

Genomic DNA samples were quantified using a Qubit fluorometer. For each sample, 100 ng of DNA was fragmented to an average size of 350 bp by ultrasonication (Covaris ME220 ultrasonicator). DNA sequencing libraries were prepared using dual-index adapters with the TruSeq Nano DNA Library Prep kit (Illumina) as per the manufacturer’s protocol. The amplified libraries were checked on a Tape Station (Agilent Technologies) and quantified by real-time PCR using the KAPA Library Quantification kit (Roche) with the QuantStudio-7flex Real-Time PCR system (Thermo Fisher Scientific). Equimolar pools of sequencing libraries were sequenced using S4 flow cells in a Novaseq 6000 sequencer (Illumina) to generate 2 x 150-bp sequencing reads for 30x genome coverage per sample.

### Extracting methylation base-called reads and QC

Sequence information was encoded in signal-level data measured with a Nanopore sequencer. The raw signal data in Fast5 format was then demultiplexed using the Guppy base-caller, and the adapters were removed using Porechop to obtain the base-called reads. The quality of the reads (in Fastq format) following adapter removal was evaluated using Fastqc (Supplementary Figure 1). The adapter-trimmed reads in Fastq format were then mapped to the human reference genome (GRCh37, patch 13, using the Minimap2 alignment program, version 2.17). The aligned reads were then used to identify the methylated sites using Nanopolish software (version 0.13.2). This tool provides a list of reads with the log-likelihood ratios (wherein a positive value is evidence that the cytosine is methylated to 5-mC, and a negative value indicates that an unreacted cytosine is present). From this file, we obtained the consolidated methylation frequency file for both the sequenced samples. Average read lengths of 5.77 kb and 12.15 kb were obtained for the ZPMetG-HV2a-1A (t0, old) and ZPMetG-HV2a-1B (t0+12, recent) respectively.

### Differential methylation analysis

The reads with methylated CpGs from the methylation frequency files of the two samples (ZPMetG-HV2a-1A and ZPMetG-HV2a-1B) were compared using bedtools (Version.v2.29.2). Reads with corresponding entries available in the two samples were taken further for differential methylation analysis.

To the above, methylated reads common to the samples, ZPMetG-HV2a-1A and ZPMetG-HV2a-1B annotations were provided from the GRCh37 GTF file.

### Annotations were provided in the following manner

- Promoter regions were defined as extending 2 kb upstream of genic regions
- TSS for genes on the “+” strand, column 2 and column 3 for genes on the “– “strand
- Genic and nongenic regions were extracted based on the tags provided in the third column of the GTF file

### Hypomethylated and hypermethylated regions

The porechopped Fastq reads for the two samples were mapped back to the reference genome GRCh37 (patch 13) using the Minimap2 program, and the aligned outputs in the binary alignment map (BAM) were then represented in BED format, which was carried out to identify the regions sequenced in both samples. The unique regions in each sample that were methylated corresponded to hypomethylated or hypermethylated regions. A region was tagged as only “hypermethylated” if there were reads available for the given region in the sample ZPMetG-HV2a-1A but not reported in the Nanopolish-generated methylation frequency file. It was tagged as only “hypomethylated” if it was reported in the Nanopolish-generated methylation frequency file for ZPMetG-HV2a-1A but no such corresponding record in ZPMetG-HV2a-1B for a region sequenced with reads was available. The gene annotations for these regions were obtained in the same manner as discussed in the previous section.

### Read mapping and variant calling for Illumina sequencing (ZPMetG-HV2a-1A and ZPMetG-HV2a-1B)

Single-nucleotide variants (SNVs) and indels were called using four different pipelines through a combination of two read mappers and two variant callers. The GRCh37 (patch 13) human genome was used as the reference genome to map the paired-end reads. The two read mappers used were BWA-MEM (version 0.7.17) and Bowtie2 (version 2.4.1). The variant-calling pipeline was implemented using GATK (version 7.3.1).

The GATK pipeline included additional read- and variant-processing steps, such as duplicate removal using Picard tools, base quality score recalibration, indel realignment, and genotyping and variant quality score recalibration using GATK, all performed according to GATK best-practice recommendations^18^. The MuTect2 tool^19^ was employed for identifying putative somatic variants, following which Snpeff (build 2017-11-24) was used to annotate the variants and make functional predictions. As described later in the Results section, variants identified using the BWA + GATK pipeline were used for all downstream analyses. Variants in the intersection of all four pipelines (two read mappers and two variant callers), in which the intersection is defined as variant calls for which the chromosome, position, reference, and alternate fields in the VCF files were identical, were confidently identified.

### Pathway mapping of genes to assess physiological implications

A compendium of genes obtained from gene set enrichment analysis (GSEA) were extracted that corresponded to tumor suppressors and oncogenes. Housekeeping genes were collated from (source). Genes associated with aging-regulating pathways were obtained from KEGG and other published work(s). KEGG pathway genes associated with different neurological and physiological disorders were extracted. For this set of genes, epigenetic changes between the two samples were examined. These epigenetic changes were measured in terms of differences in methylation frequencies for ZPMetG-HV2a-1A and ZPMetG-HV2a-1B. The frequency of methylation is given by the ratio of the number of methylated reads in a gene region (**m**_**r**_) and the total number of reads detected in the gene region (**T**_r_).

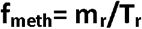

The difference in methylation frequency (**Δf**_**meth**_) between the ZPMetG-HV2a-1B (t_0_+12) and ZPMetG-HV2a-1B (t_0_ years) was obtained by subtracting the methylation frequency for t_0_ from that of t_0_+12.

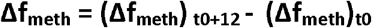

The degree of methylation was primarily classified as hypomethylated, hemi-hypomethylated, hemi-hypermethylated, and hypermethylated based on the int **Δf**_**meth**_ intervals. Here, the term “background” was also defined as corresponding to low-confidence scores of CpG methylation, namely, –0.2 ≤ **Δf**_**meth**_ < +0.2. Hypomethylation was defined as a gradient of hypomethylation (**Δf**_**meth**_ < 0.6) and hemi-hypomethylation (–0.6 ≤ **Δf**_**meth**_ < –0.2). Hypermethylation was defined as a gradient of hypermethylation (**Δf**_**meth**_ > than 0.6) and hemi-hypermethylation (0.2 ≤ **Δf**_**meth**_ < 0.6). The weighted difference in methylation frequency (**Δf**_**meth_weighted**_) for each read within a gene region was obtained by factoring in the number of CpG islands (**cpg**_**num**_) detected for that gene region. Thus, the weighted methylated frequency was obtained as a product of the difference in methylation frequency and the number of CpG reads in the gene region.

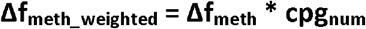

The normalized weighted methylation frequency score (**F**_**meth_norm**_) per degree of methylation was obtained by normalizing the summed **Δf**_**meth_weighted**_ score with the total number of reads (T_r_).

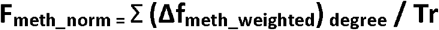

The normalized weighted difference in methylation frequency score was assigned to a specific gene and used in downstream analysis.

### Mutational signature analysis

The non-negative matrix factorization (NMF) method described by Alexandrov et.al^20^., was used to detect mutational signatures for disease associated KEGG gene sets in ZPMet-Hv2a-1A and ZPMet-Hv2a-1A WGS. To determine the contribution of each signature in a sample, we used flexible functions in the MutationalPatterns^21^ routines, available in the R package, to assign the mutations in each sample to the signatures identified by NMF.

## Results

### Analysis of individual methylome modifications over time

To account for differences in coverage depth, we extracted the nucleotide ranges for which base-called entries were present in the methylation frequency files for both samples. This enabled identifying any distinctive increase or decrease in methylation between the two temporally separated samples. These comparable nucleotide ranges extracted from the methylation base-called regions accounted for 18,758,642 regions, which covered 98% and 84% of reported regions in fastq files of ZPMetG-HV2a-1A (old) and ZPMetG-HV2a-1B (recent) respectively. The overall analysis pipeline is detailed in Figure 1A. We examined the overall distribution in methylation frequency between ZPMetG-HV2a-1A (old) and ZPMetG-HV2a-1B (recent) We applied a nonparametric statistical test (Mann–Whitney U test) and observed a statistically significant difference between the two sample distributions (Figure 1B). The chromosome-wise distribution of methylated frequencies in CpG dinucleotides, which was also examined to discern changes in methylation between the samples, showed similar profiles for genic and intergenic regions. Notable exceptions were observed for chromosomes 21, 22, and X for genic regions (Supplementary Figure 2). The median of the distribution is given for each chromosome within the entire frequency distribution (Figure 1B).

**Figure 1.**
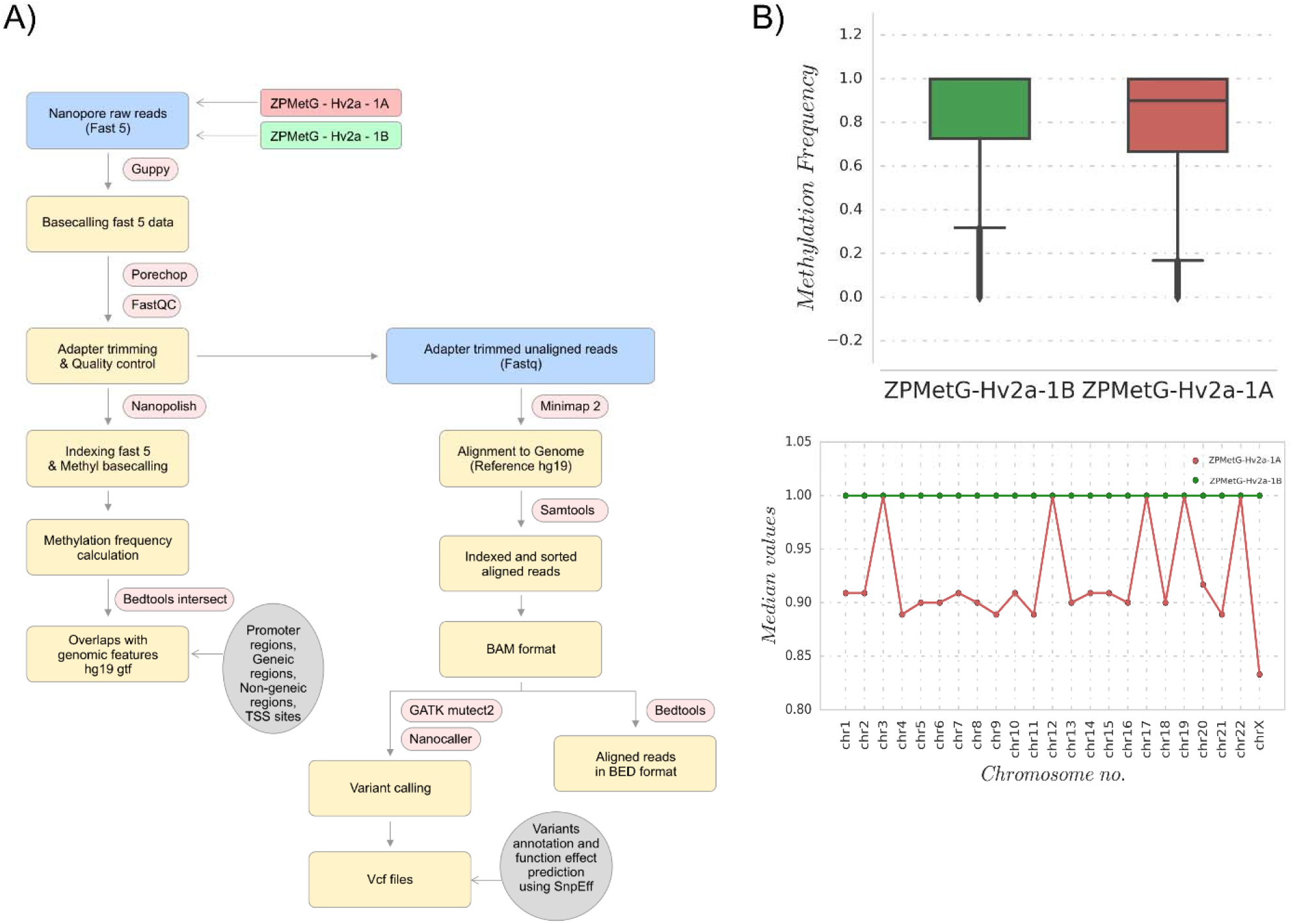
**(A)** Workflow for the analysis of individual methylome and mutational (SNP) variations for two samples from the same individual collected 12 years apart. **(B, upper)** Methylation frequency distributions for comparing base-called regions, shown for ZPMetG-Hv2a-1B (recent, green) and ZPMetG-Hv2a-1A (old, red). On the right is shown a histogram demonstrating the distribution to be non-Gaussian. **(B, lower)** Median values of the chromosome-wise methylation frequency distribution for ZPMetG-HV2a-1A (old, in red) and ZPMetG-HV2a-1B (recent, in green).

**Figure 2.**
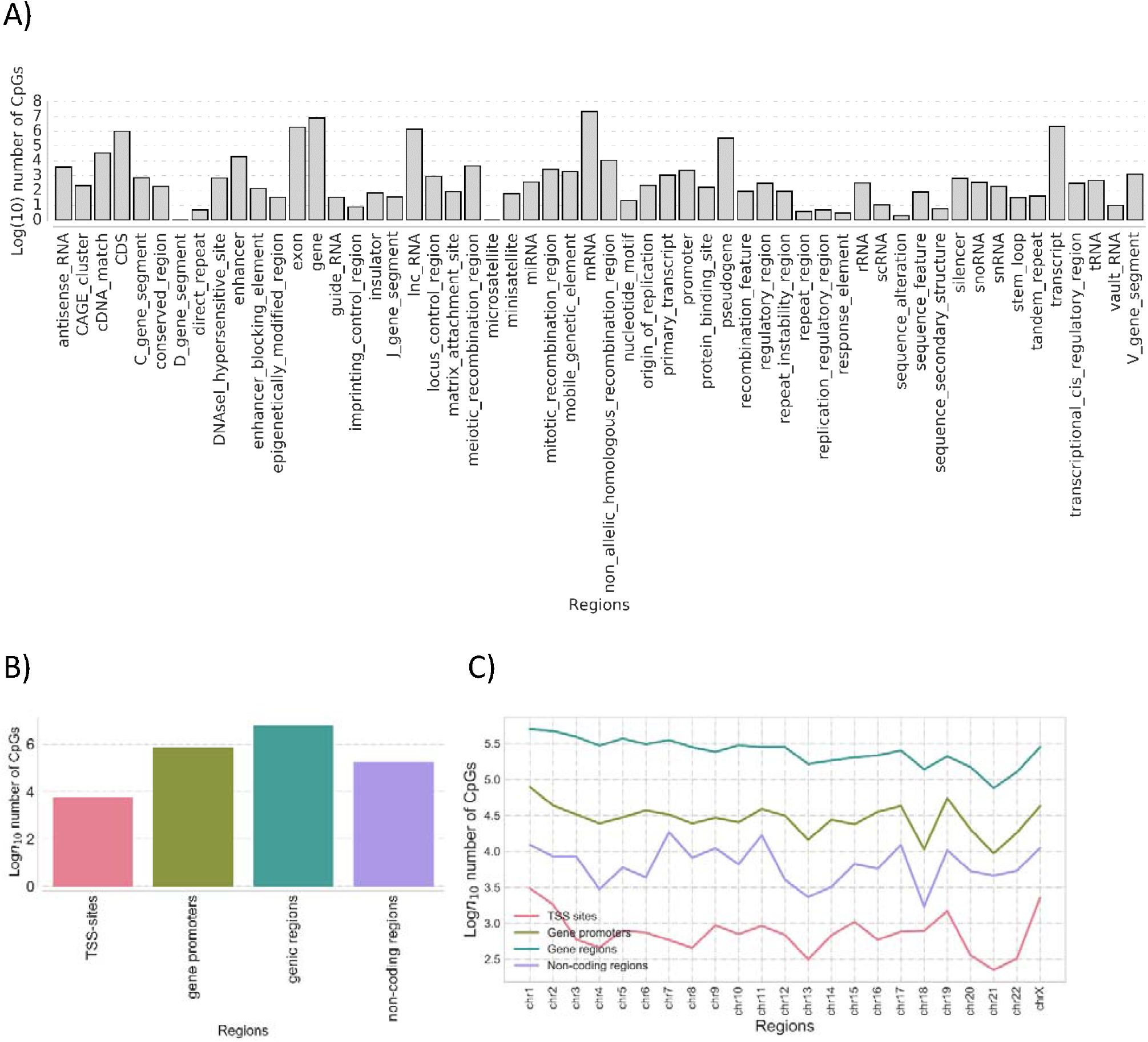
**(A)** The distribution and classification of methylation imprints (CpG) based on genome architecture and classifiers. **(B)** The classification of common methylation imprints (CpG) based on genomic characterization. **(C)** Chromosome-wise distribution of CpGs common to samples ZPMetG-Hv2a-1 and ZPMetG-Hv2a-2 classified based on their distribution across genomic motifs.

### Functional motif-based characterization of CpG’s common to ZPMetG-Hv2a-1 and ZPMetG-Hv2a-1

The methylation base-called regions compared between ZPMetG-HV2a-1A (old) and ZPMetG-HV2a-1B (recent) were mapped to GRCH37 (human reference genome from NCBI). Based on the tag “gene”, we extracted putative genic regions, while other regions (excluding even CDS, mRNA, and tRNA) were extracted to examine the non-genic regions (Figure 2A). A majority of the CpGs were found in the genic and gene promoter regions (Figure 2B). The distribution across chromosomes for genic and non-genic regions in the two samples indicated that a majority of the CpGs occurred within genic regions and were rarest within TSS regions. The distribution of CpGs was similar across all chromosomes for the genic regions compared with the distribution for TSS, non-genic, and genic promoter regions, which displayed a staggered distribution across chromosomes (Figure 2C).

### Prioritization of gene families

Characterization of the methylated regions indicated that 26,019 genes are found in the regions methylated in ZPMetG-HV2a-1B (recent) compared to ZPMetG-HV2a-1A (old) and selective filtering and prioritization of the 26,019 genes (normalized score) based on a curated dataset of 3050 gene families from the GSEA database^22^, 168 from the Epigenetic Modifier Gene database^23^, 2833 housekeeping genes from HRT Atlas v1.0^24^, and 3527 pathway- and disease-specific genes from the KEGG database^25^ yielded 5214 unique genes. Followers of the

Zoroastrian faith consider smoking a taboo, resulting in generations of the community that have refrained from smoking. To address the epigenetic changes associated with smoking-related genes and their relevance to the non-smoking Parsi community, we also curated a list of 44 documented genes that were differentially methylated in smokers. Taken together, our total gene list was comprised of 5258 genes (Supplementary Table.2).

### Hierarchical clustering of differentially methylated regions in ZPMetG-Hv2a-1B compared to ZPMetG-Hv2a-1A

We performed hierarchical clustering of the differentially methylated genes by first obtaining the frequency of CpG methylation across the entire gene length. Based on the weighted score for CpG methylation (Materials and Methods), the CpG frequencies across genes were classified as hypo-, hemi-hypo-, hemi-hyper-, and hypermethylated regions. The weighted score common across all genes was defined as the background, which served as an internal control.

The hierarchical clustering of row-wise Z score of common differentially methylated gene regions (Supplementary Table 3) in ZPMetG-Hv2a-1B identified a cluster of 103 genes that was significantly hypermethylated and 307 genes that were hypomethylated (Figure 3A, Supplementary Table 7). GSEA-based enrichment indicated an enrichment in genes coding for transcription factors and protein kinases in the hyper- and hypomethylated clusters. Among the hyper-methylated genes, homeodomain proteins were highly represented, while among the hypomethylated genes, cell differentiation markers, tumor suppressors, and translocated cancer genes predominated by comparison (Figure 3B, Supplementary Figure 3A)

**Figure 3.**
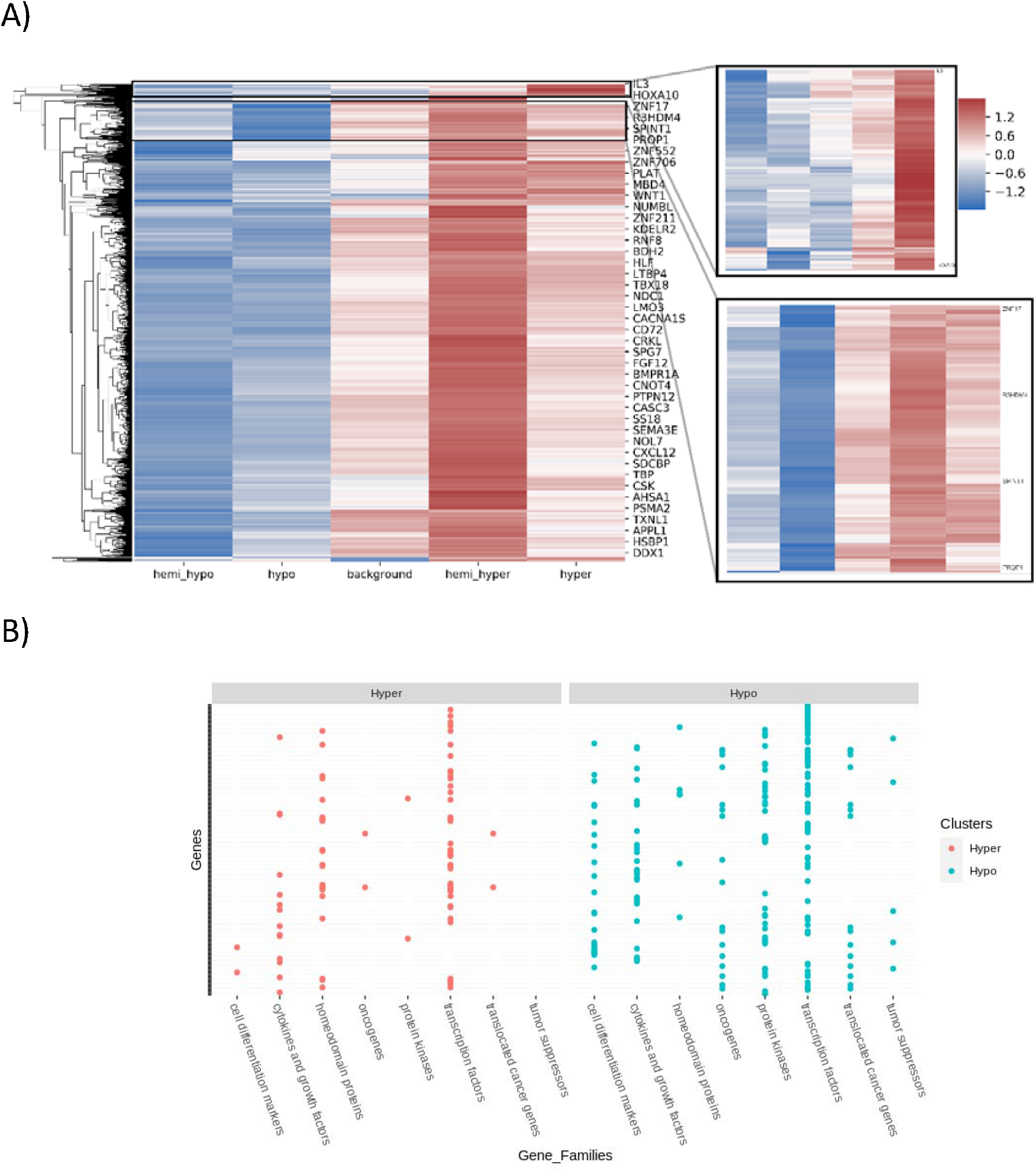
**(A)** Hierarchical clustering of row-wise Z scores of common differentially methylated gene regions in sample s2. Clusters of hypermethylated (inlay figure; right, top) and hypomethylated (inlay figure; right, bottom) genes. **(B)** GSEA-based classification of hyper- and hypomethylated clusters.

PANTHER-based gene annotation classified these genes based on their molecular and protein functions (Supplementary Figure 3B). Hypermethylated and hypomethylated clusters varied in their functional classification with the presence of genes that function to increase transcription regulator activity (Supplementary Figure 3B).

### Characterization of variants unique to ZPMetG-Hv2a-1B compared with ZPMetG-Hv2a-1A

While dynamic methylation events constitute a major mechanism of epigenetic modification, variants in the form of SNPs, in addition to dynamic methylation events, are known to be critical to epigenome function through the modification of genomic information. To this end, we employed the GATK variant characterization pipeline to identify variants specific to ZPMetG-Hv2a-1B viz., which are variants that have accumulated in the individual because of aging. We identified 4,58,148 variants corresponding to 24,948 unique genes.

We next sought to identify genes common to both dynamic methylated regions (CpGs) and variants specific toZPMetG-Hv2a-1B. We found a direct correlation between gene length, CpG count, and variant count. This association is specifically significant for the correlation between variants and CpG (Pearson correlation coefficient, r=0.86, Figure 4A, Supplementary Figure 4A, B). Analysis of the 5258 ZPMetG-Hv2a-1B-specific differentially methylated genes post disease/GSEA prioritization filter for somatic variants yielded 4409 differentially methylated genes (Supplementary Table 5) that are specific to ZPMetG-HV2a-1B (recent) and that harbour somatic variants (Supplementary Figure 4C).

**Figure 4.**
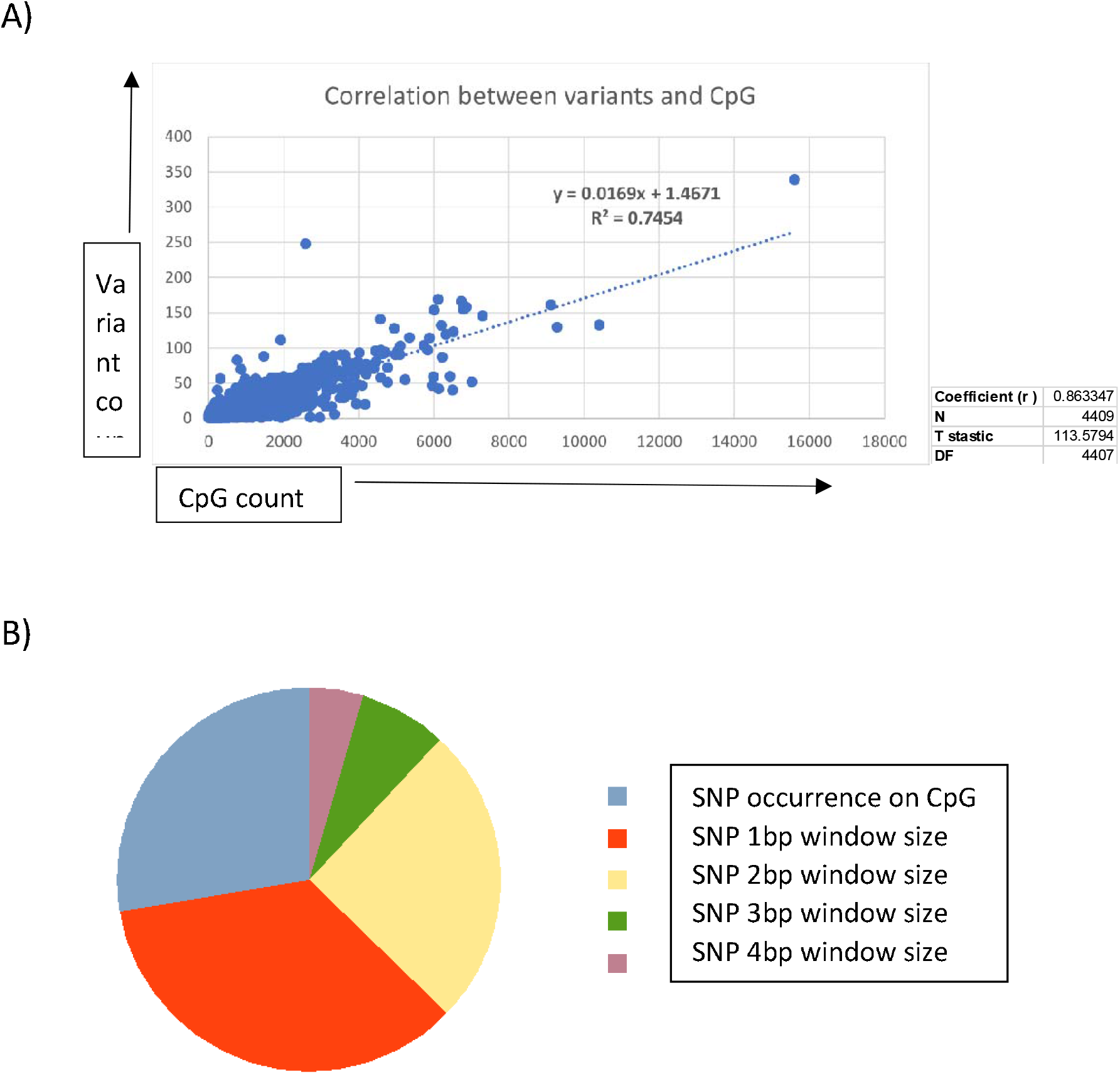
**A)** Correlation between CpG and variant counts in ZPMetG-Hv2a-1B. The Pearson correlation coefficient, r, is displayed at the side of the plot. **B)** Distribution of ZPMetG-HV2a-1B (recent) specific variants across differentially methylated regions specific to ZPMetG-HV2a-1B (recent).

The location of variants with respect to their occurrence in CpG regions is crucial to the resulting effect on epigenomic imprinting in the form of methylation changes. Variants that occur exactly at the CpG site may modulate the resulting methylation, thereby effecting methylation changes. In this context, we classified ZPMetG-HV2a-1B variants based on their locus of incidence within the CpG region. We classified variant occurrence as those that occur exactly within the CpG region and variants that occur within a distance window of 1–4 bp on either side of the CpG region (±4 bp). Our analysis indicated 242 variants classified as “modifier” variants occurring within the CpG region of 177 differentially methylated genes specific to ZPMetG-Hv2a-1B (Figure 4B, Supplementary Table 6). A fraction of variants (67/242 or 27.68%) occurred exactly at the CpG site and 175/242 variants (72%) occurred in the CpG region at a locus window of ±4 bp (Fig. 4B). A vast majority of classified variants occurred on Chr. 1, with most variants occurring within a ±4-bp window across all chromosomes except for Chr. 4, in which more variants are observed exactly at the CpG site compared with the window size (Supplementary Figure 5A). The number of variants within genes varied from 1–7 variants (Supplementary Figure 5B). Characterization of the pathway association and interaction networks differed between the genes harbouring variants at the CpG site compared with genes with variants in the CpG region (Supplementary Figures 6 and 7).

### Interplay of the genome and epigenome in the context of aging and disease etiology

The presence of genome variants can affect epigenetic modification. We therefore sought to understand the biological processes of the genes in our study that carried both methylation changes and harbored variants in the CpG regions. To this end, we compared 412 genes with significant hypermethylation (108 genes) and hypomethylation (304 genes) changes (Supplementary Table 7) for gene-specific variants within the CpG regions. Our analysis yielded a critical cluster of 10 genes that were significantly methylated (hyper- or hypo-) and have variants at the CpG site or within the ±4 bp CpG-region window (Figure 5A, Supplementary Table 8). Only two genes (*CASP8* and *PCGF3*) were significantly hypomethylated and carried variants at the CpG site, whereas both significantly hyper/hypo-methylated genes carried variants in the CpG window region (3 hypermethylated genes and 6 hypomethylated genes). The vast majority of genes were critical housekeeping genes, followed by transcription factors, protein kinases, and genes implicated in neurodegenerative diseases (Figure 5B).

**Figure 5.**
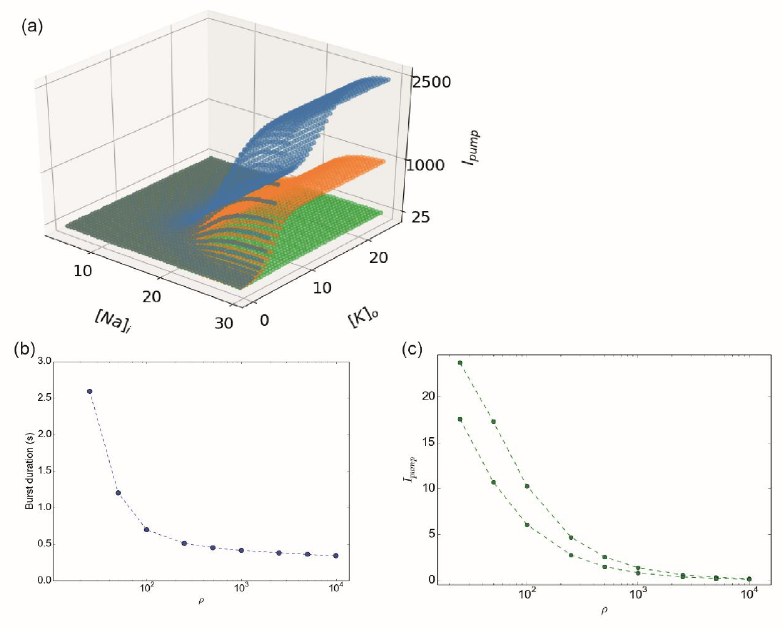
**A)** Distribution of CpG variants (exact, inexact) across significantly hyper- and hypomethylated genes in ZPMetG-Hv2a-1B **B)** Categorization of 10 critical genes based on biological activity. **C)** Network analysis using a KEGG-based disease-enrichment module for 10 critical and 1000 high-priority variants. **D)** Reactome analysis of biologically relevant pathways of 10 critical and 1000 high-priority variants.

To identify the relevance of these 10 genes in pathways related to disease and homeostasis, we clustered the 10 genes with 1000 genes that harboured variants and CpGs (Supplementary Table 9, based on the normalized cumulative CpG.Variant score). Using Network analyst^26^-based clustering and a KEGG-based disease enrichment module showed that the majority of the clustered genes were implicated in cell proliferative pathways regulating cancer; cancer subtypes; neurodegenerative diseases, such as Parkinson’s disease, Alzheimer’s disease, Huntington’s disease; and pathways implicated in cellular senescence and longevity-regulating gene clusters (Figure 5C, Supplementary Table 10). Reactome analysis of the biological diversity of gene function showed the association of the clustered genes with pathways involved in the immune system, DNA repair, DNA transcriptional activity, the cell cycle, and disease-related pathways (Figure.5D).

### Mutational signatures associated with aging in the ZPMetG-Hv2a-1B across all genes and CpG-specific regions of functionally relevant genes

We next proceeded to categorise mutational signatures that are characteristic combinations of mutation types arising from specific mutagenesis processes that involve alterations related to DNA replication, repair, and environmental exposure. Our analysis of showed the presence of 110,616 mutations, with a majority associated with C>T transitions, followed by T>C transitions. The same analysis of 4409 prioritized genes that showed differential methylation and variants specific to indicate a similar signature for C>T transitions, followed by an increase in C>A and T>A mutations, while T>C mutations were reduced compared with the cumulative variants in ZPMetG-Hv2a-1B (Figure 6A). Further categorization of the approach using gene sets that consist of epigenetic modifiers, disease-specific lists, and smoking-related genes (sensitive to environmental exposure) showed a high prevalence of C>T transitions, especially at the CpG sites for gene sets regulating Alzheimer’s and Parkinson’s disease pathways, while smoking-associated genes had a decreased incidence of C>T transitions at CpG sites but an increase in overall C>T transitions (Figure 6B). Analysis of the complete 96 mutational signatures indicated the prevalence of signatures for Parkinson’s, Alzheimer’s, and Huntington’s diseases, while mutational signatures for smoking-associated genes were not highly represented (Figure 6C).

**Figure 6.**
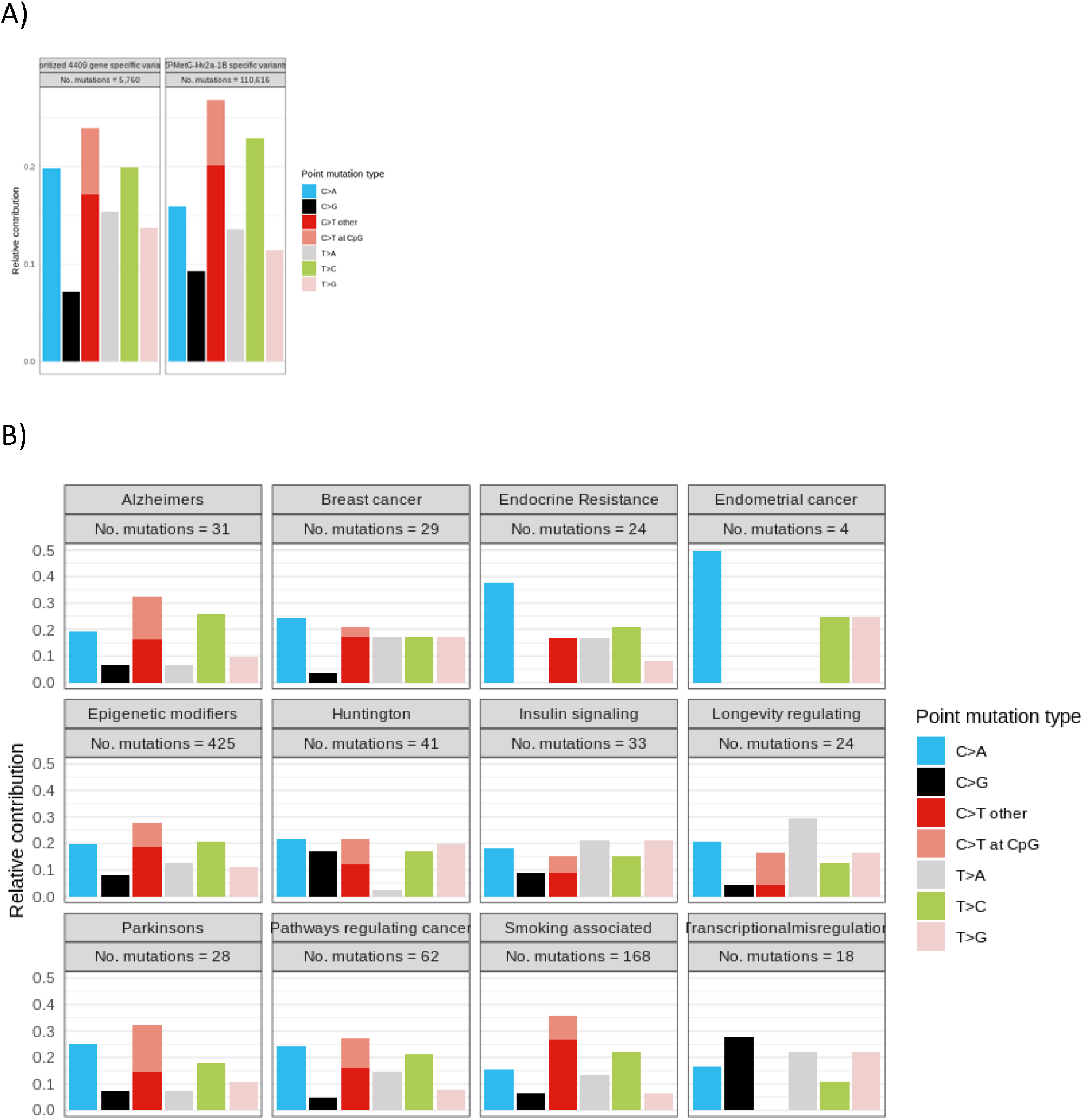
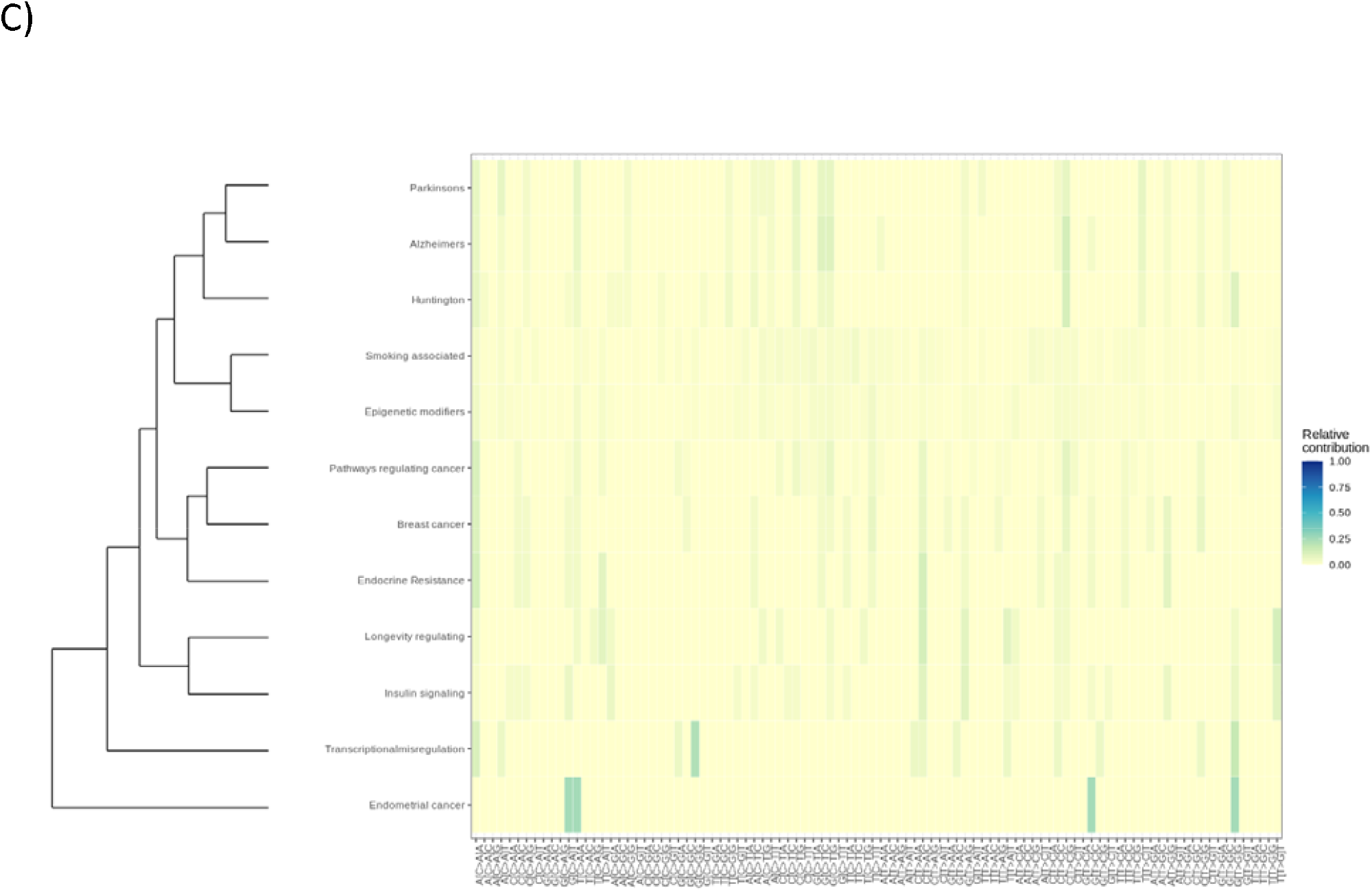
**A)** Distribution of mutational signatures across the genome and specific to CpG regions in 4409 prioritized genes and cumulative variants across ZPMetG-Hv2a-1B. **B)** Mutational signatures indicating transitions and transversions specific to ZPMetG-Hv2a-1B across different disease categories. **C)** Analysis of the relative contribution of the complete repertoire of mutational signatures (96) specific to ZPMetG-Hv2a-1B.

## Discussion

Aging is an important contributor to chronic diseases. Epigenetic modifications in the form of methylation of CpG dinucleotides play a crucial role in regulating physiological processes, which vary as age progresses. Differential methylation at CpG sites between younger and older subjects has been shown to be associated with genes, including MTOR, ULK1, ADCY6, IGF1R, CREB5, and RELA, that are linked to metabolic traits^27^, and CREB5, RELA, and ULK1 have been statistically associated with age. We found 5258 genes to be significantly altered in terms of methylation differences over time in an individual assessed after 12 years. DNAm levels of several CpG sites located within genes involved in longevity-regulating pathways can be examined for DNAm in metabolic alterations associated with age progression.

Along with epigenetic modifications, variants at the CpG sites have been shown to adversely affect gene function and activity, especially for disease-associated traits^28^. Many studies have reported that CpG–SNP interactions are associated with different diseases, such as type 2 diabetes, breast cancer, coronary heart disease, and psychosis, that show a clear interaction between genetic (SNPs) and epigenetic (DNA methylation) regulation. The introduction or removal of CpG dinucleotides (possible sites of DNA methylation associated with the environment) has been suggested as a potential mechanism through which SNPs influence gene transcription and expression via epigenetics. Our analysis showed the presence of 242 variants, classified as “modifier” variants, occurring within the CpG region of 177 differentially methylated genes specific to ZPMetG-Hv2a-1B. These are genes involved in longevity-regulating pathways, cellular proliferation, and transcriptional regulation.

Recent reports using tissue-specific methylation data describe a strong association between C>T mutations and methylation at CpG dinucleotides in many cancer types, driving patterns of mutation formation throughout the genome. In our analysis, the variants occurring at the CpG sites were C>T transitions that occur within two genes, *CASP8* and *PCGF3*. Incidentally, in our analysis *PCGF3* displays transitions not only at the CpG site but also within the CpG region. *PCGF3/5* mainly functions as a transcriptional activator, driving expression of many genes involved in mesoderm differentiation, and *PCGF3/5* is essential for regulating global levels of the histone modifier H2AK119ub1 in embryonic stem cells^29^. *CASP8* methylation has been shown to be relevant in cancers and may function as a tumor-suppressor gene in neuroendocrine lung tumors^30^. Our study points to a crucial interplay between CpG methylation levels and variant incidence. It is therefore critical to extend the study to other individuals to understand the diversity of mutational signatures that accrue in CpG regions of functionally important genes to further dissect the role of epigenome–genome interactions in homeostasis, disease, and aging.

Mutational signatures associated with aging and cancer subtypes have been studied and indicate a strong correlation towards cytosine deamination at CpG sites, which results in an increase in the frequency of C>T transitions with age of diagnosis, as age allows more time for deamination events, leading to an accumulation of their effects^31^. In line with these observations, we found that C>T transitions constitute most of the mutational changes in ZPMetG-Hv2a-1B, indicating their accumulation over age. Specifically, we found an increase in frequency of the same transitions at CpG sites in genes that participate in Alzheimer’s and Parkinson’s disease pathways, but their relative contribution is decreased in smoking-associated genes. Further analysis of the same signatures in a larger cohort corresponding to disease and smoking status will be critical in identifying the combinatorial effect of mutational signatures associated with epigenome modifications in the context of aging.

By means of personal methylome analysis, our study furthers understanding of the interactions between the epigenome and genome in the form of CpG–gene variant interactions. This approach promises to be helpful in identifying significant genomic loci that reflect the effects of methylation modifications in combination with genetic variations that accrue in an individual, thereby aiding in identifying associations with longevity and associated traits, such as the timing of disease onset.

