## Supplementary Figures for "Dynamic methylome modification is associated with mutational signatures in aging and the etiology of disease"

#### Supplementary Figure 1

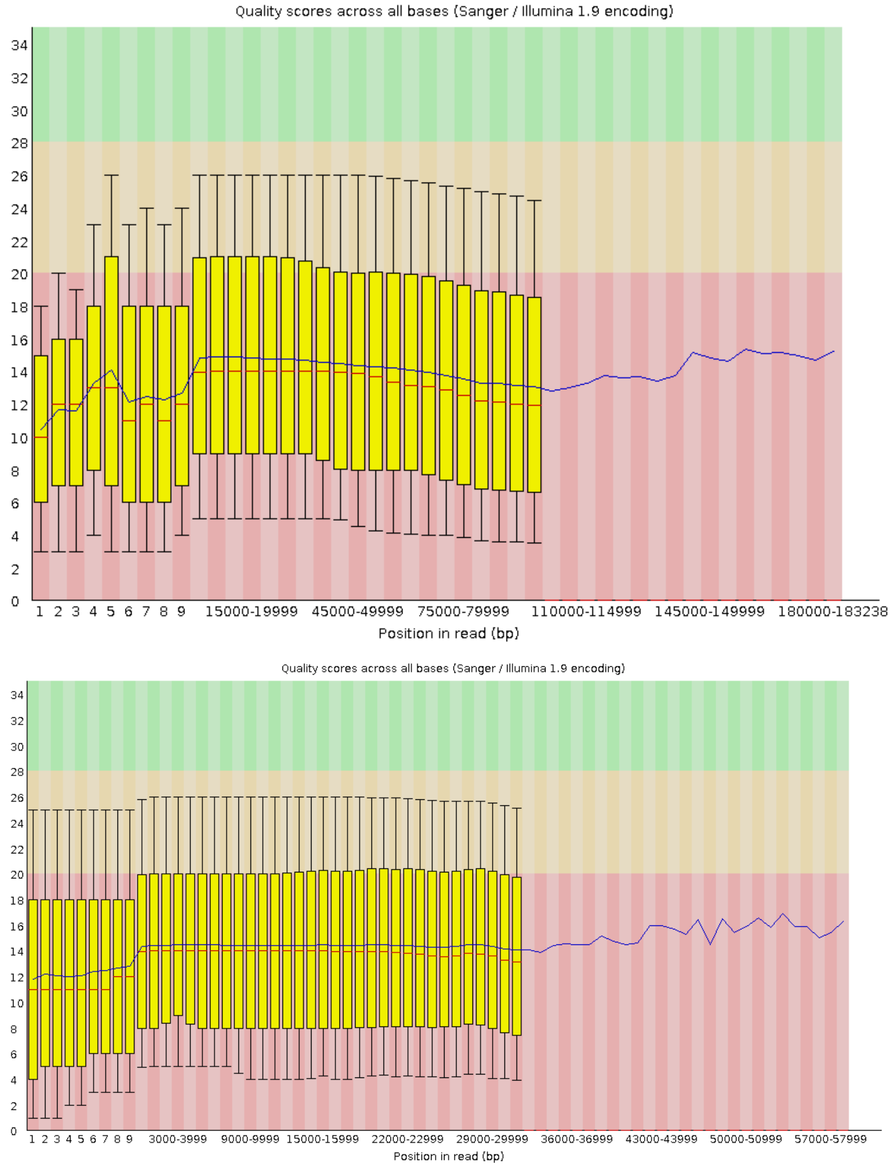


**Supplementary Figure 1.** QC scores (FastQC) of s1 and s2 following adapter trimming.

#### Supplementary Figure 2

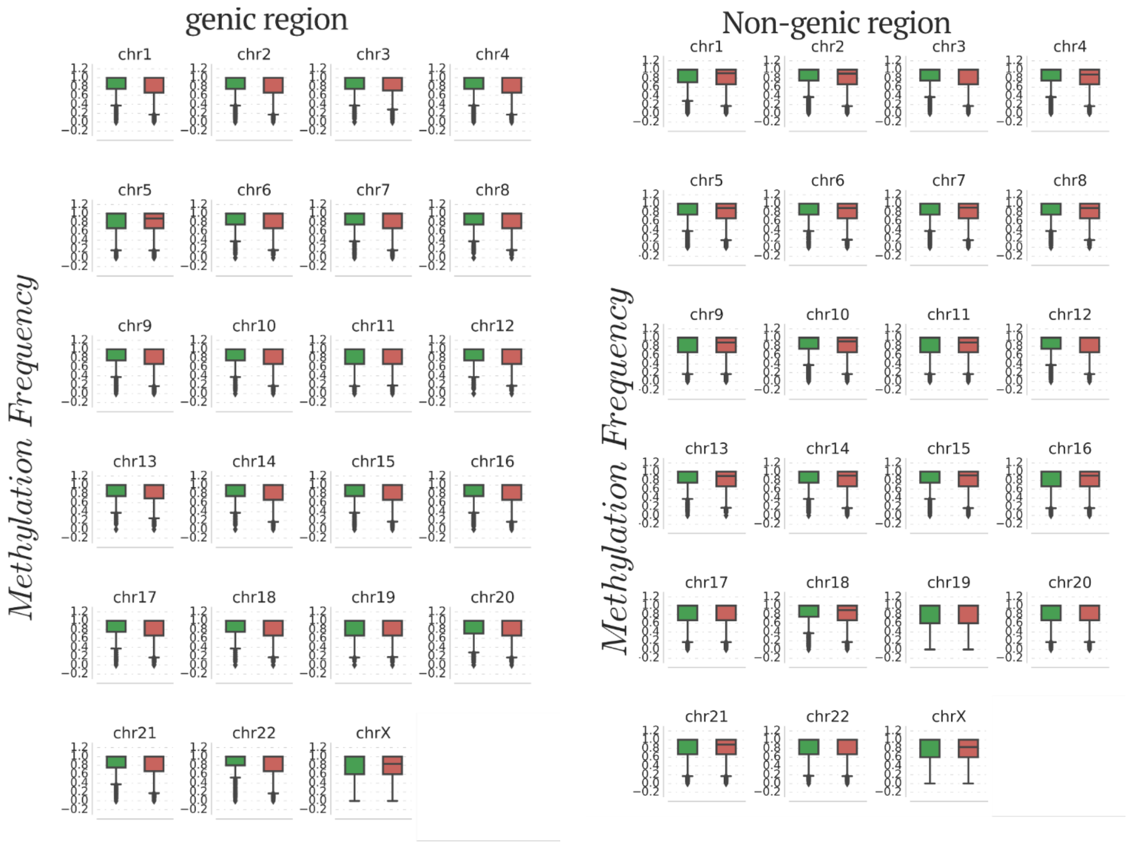


**Supplementary Figure 2.** Chromosome-wise distribution of methylation frequency across genic and nongenic regions for ZPMetG-Hv2a-1B.

### Supplementary Figure 3

A

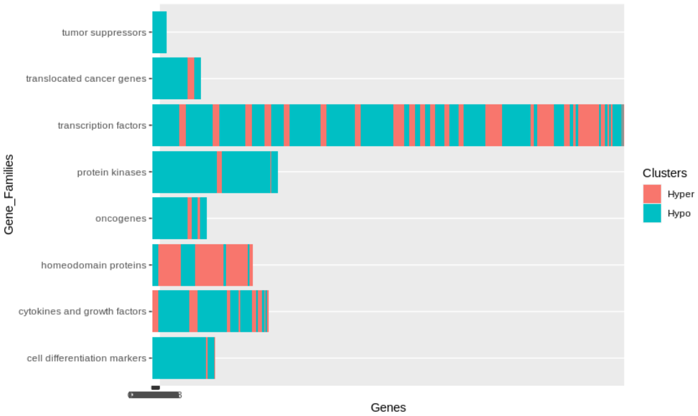


B

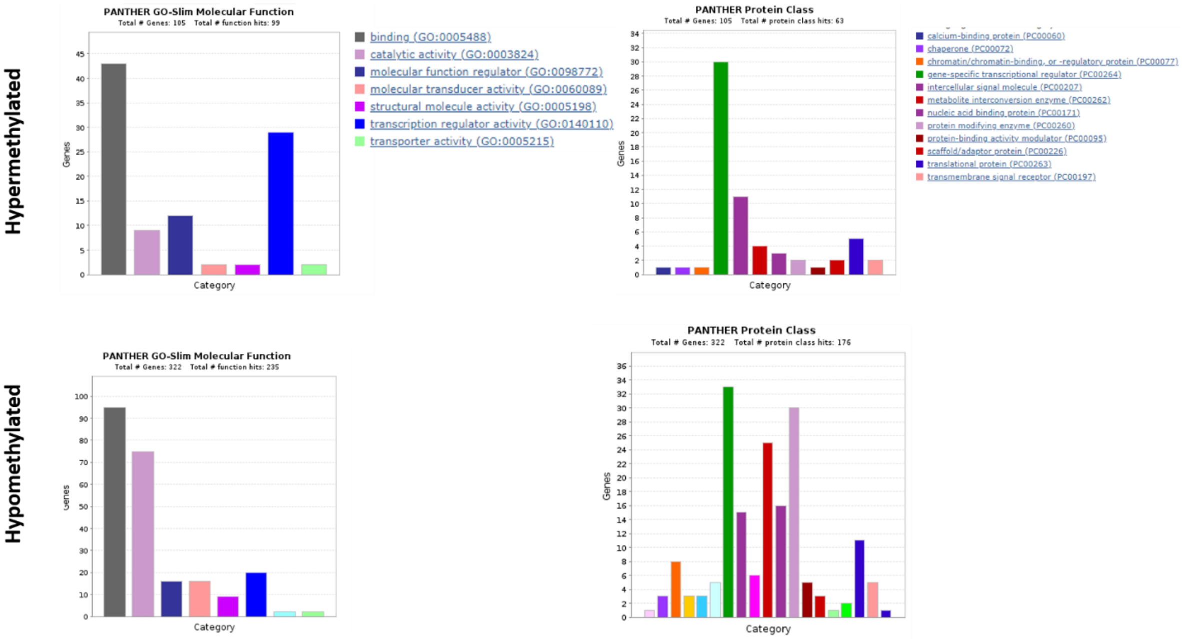


**Supplementary Figure 3.** Molecular and protein classification of significantly hyper- and hypomethylated genes. **A)** GSEA-based categorization of selected hyper- and hypomethylated genes specific to ZPMetG-Hv2a-1B. **B)** PANTHER-based annotation of hyper- and hypomethylated genes based on biological and molecular functions.

### Supplementary Figure 4

A)

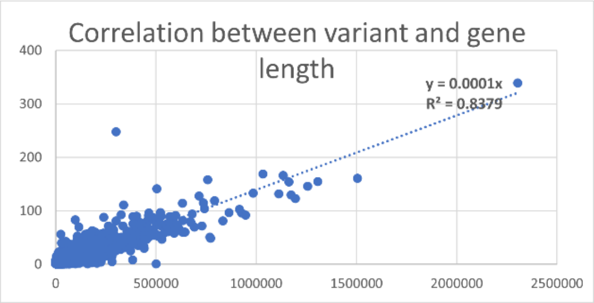


B)

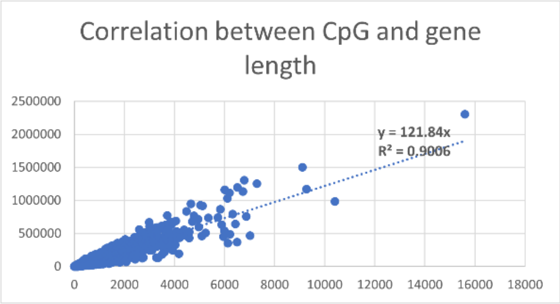


C)

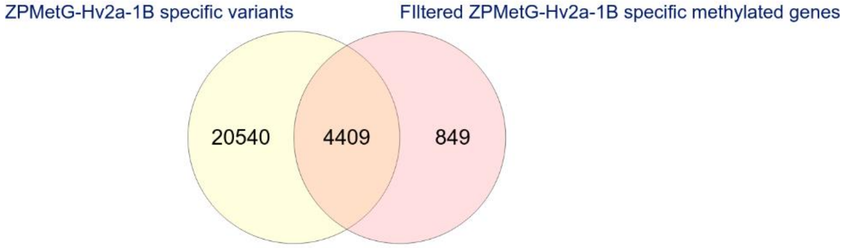


**Supplementary Figure 4. A)** Correlation between the number of variants and gene length. **B)** Correlation between the number of CpGs and gene length. **C)** Intersection of s2-specific gene variants and prioritized methylated genes specific to ZPMetG-Hv2a-1B.

#### Supplementary Figure 5

A

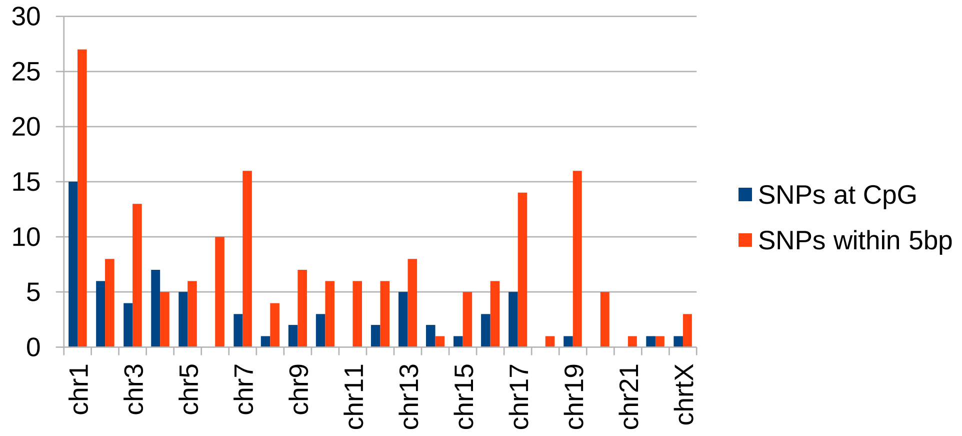


B

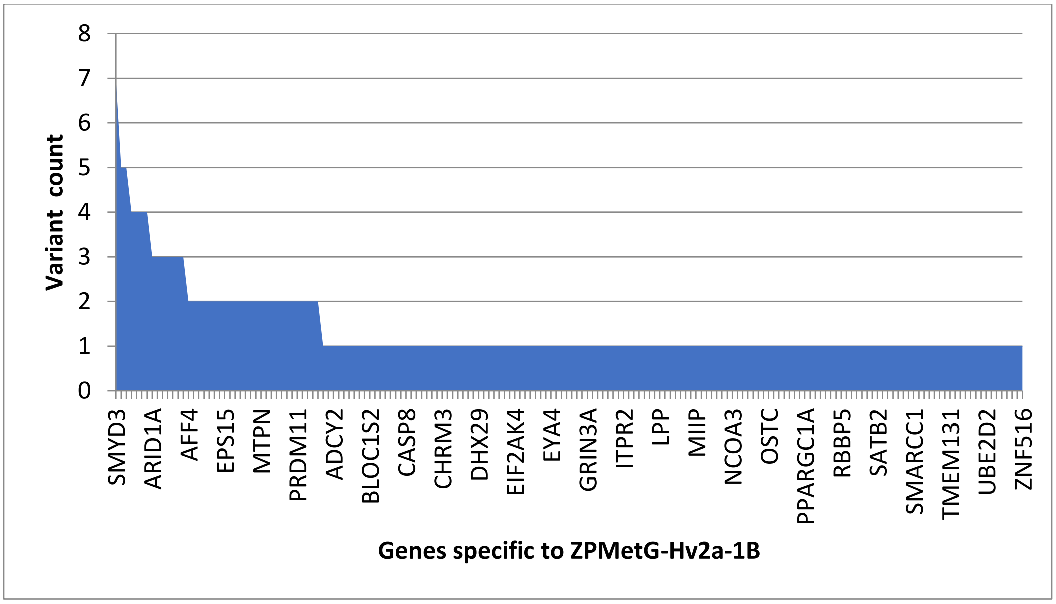


**Supplementary Figure 5.** Classification and variant information specific to ZPMetG-Hv2a-1B. **A)** Chromosome-wise distribution of variants across the CpG region of differentially methylated genes specific to ZPMetG-Hv2a-1B. **B)** Variant count per gene for genes specific to ZPMetG-Hv2a-1B.

### Supplementary Figure 6

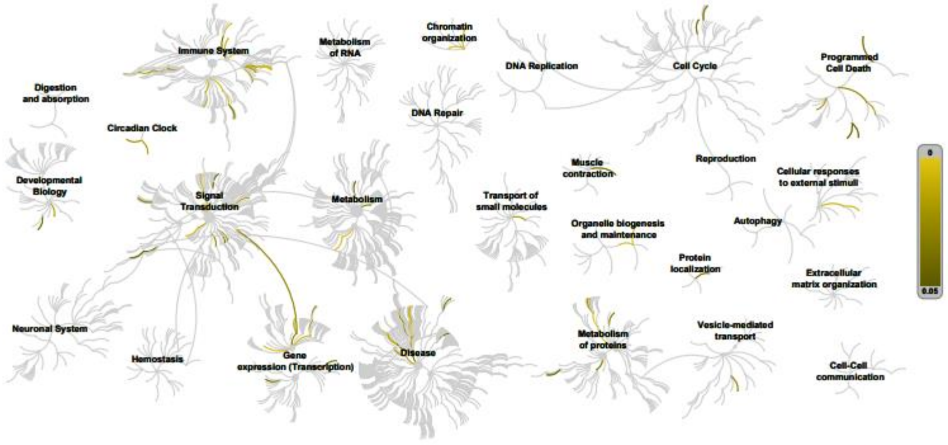


| Pathway name | Entities | | | | Reactions | |
|---|---|---|---|---|---|---|
| | found | ratio | p-value | FDR* | found | ratio |
| Circadian Clock | 6 / 104 | 0.007 | 9.35e-06 | 0.004 | 48 / 59 | 0.004 |
| SUMO E3 ligases SUMOylate target proteins | 6 / 183 | 0.012 | 2.09e-04 | 0.037 | 10 / 131 | 0.01 |
| SUMOylation | 6 / 192 | 0.013 | 2.70e-04 | 0.037 | 10 / 140 | 0.011 |
| Notch-HLH transcription pathway | 3 / 28 | 0.002 | 3.14e-04 | 0.037 | 1 / 2 | 1.51e-04 |
| Signaling by NOTCH1 | 4 / 84 | 0.006 | 6.45e-04 | 0.048 | 10 / 39 | 0.003 |
| Transcriptional activation of mitochondrial biogenesis | 4 / 87 | 0.006 | 7.35e-04 | 0.048 | 12 / 32 | 0.002 |
| Erythropoietin activates STAT5 | 2 / 9 | 6.11e-04 | 8.29e-04 | 0.048 | 3 / 3 | 2.27e-04 |
| STAT5 Activation | 2 / 9 | 6.11e-04 | 8.29e-04 | 0.048 | 3 / 3 | 2.27e-04 |
| PPARA activates gene expression | 5 / 174 | 0.012 | 0.001 | 0.052 | 41 / 41 | 0.003 |
| Chromatin organization | 6 / 260 | 0.018 | 0.001 | 0.052 | 12 / 85 | 0.006 |
| Chromatin modifying enzymes | 6 / 260 | 0.018 | 0.001 | 0.052 | 12 / 85 | 0.006 |
| Regulation of lipid metabolism by PPARalpha | 5 / 176 | 0.012 | 0.001 | 0.052 | 43 / 44 | 0.003 |
| Interleukin-21 signaling | 2 / 12 | 8.15e-04 | 0.001 | 0.052 | 2 / 5 | 3.78e-04 |
| PKMTs methylate histone lysines | 3 / 49 | 0.003 | 0.002 | 0.052 | 7 / 22 | 0.002 |
| Signaling by Leptin | 2 / 13 | 8.83e-04 | 0.002 | 0.053 | 5 / 19 | 0.001 |
| CLEC7A (Dectin-1) signaling | 4 / 120 | 0.008 | 0.002 | 0.059 | 8 / 45 | 0.003 |
| NOTCH1 Intracellular Domain Regulates Transcription | 3 / 57 | 0.004 | 0.002 | 0.059 | 1 / 18 | 0.001 |
| C-type lectin receptors (CLRs) | 5 / 203 | 0.014 | 0.003 | 0.059 | 9 / 68 | 0.005 |
| Interleukin-9 signaling | 2 / 16 | 0.001 | 0.003 | 0.059 | 4 / 13 | 9.83e-04 |
| Interleukin-15 signaling | 2 / 16 | 0.001 | 0.003 | 0.059 | 5 / 17 | 0.001 |
| Mitochondrial biogenesis | 4 / 127 | 0.009 | 0.003 | 0.059 | 14 / 36 | 0.003 |
| SUMOylation of chromatin organization proteins | 3 / 62 | 0.004 | 0.003 | 0.059 | 5 / 15 | 0.001 |
| Prolactin receptor signaling | 2 / 18 | 0.001 | 0.003 | 0.059 | 2 / 14 | 0.001 |
| TICAM1, RIP1-mediated IKK complex recruitment | 2 / 19 | 0.001 | 0.004 | 0.059 | 1 / 3 | 2.27e-04 |
| Interleukin-2 signaling | 2 / 19 | 0.001 | 0.004 | 0.059 | 5 / 19 | 0.001 |

* False Discovery Rate

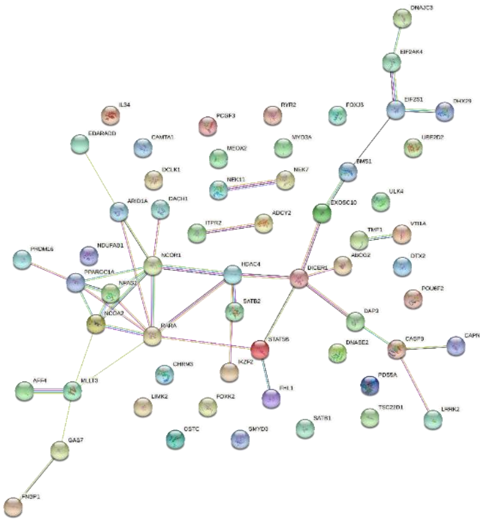

**Supplementary Figure 6.** Biochemical targets (Reactome) and pathway association (STRING) of ZPMetG-Hv2a-1B (recent) variants occurring at the CpG site (exact).

### Supplementary Figure 7

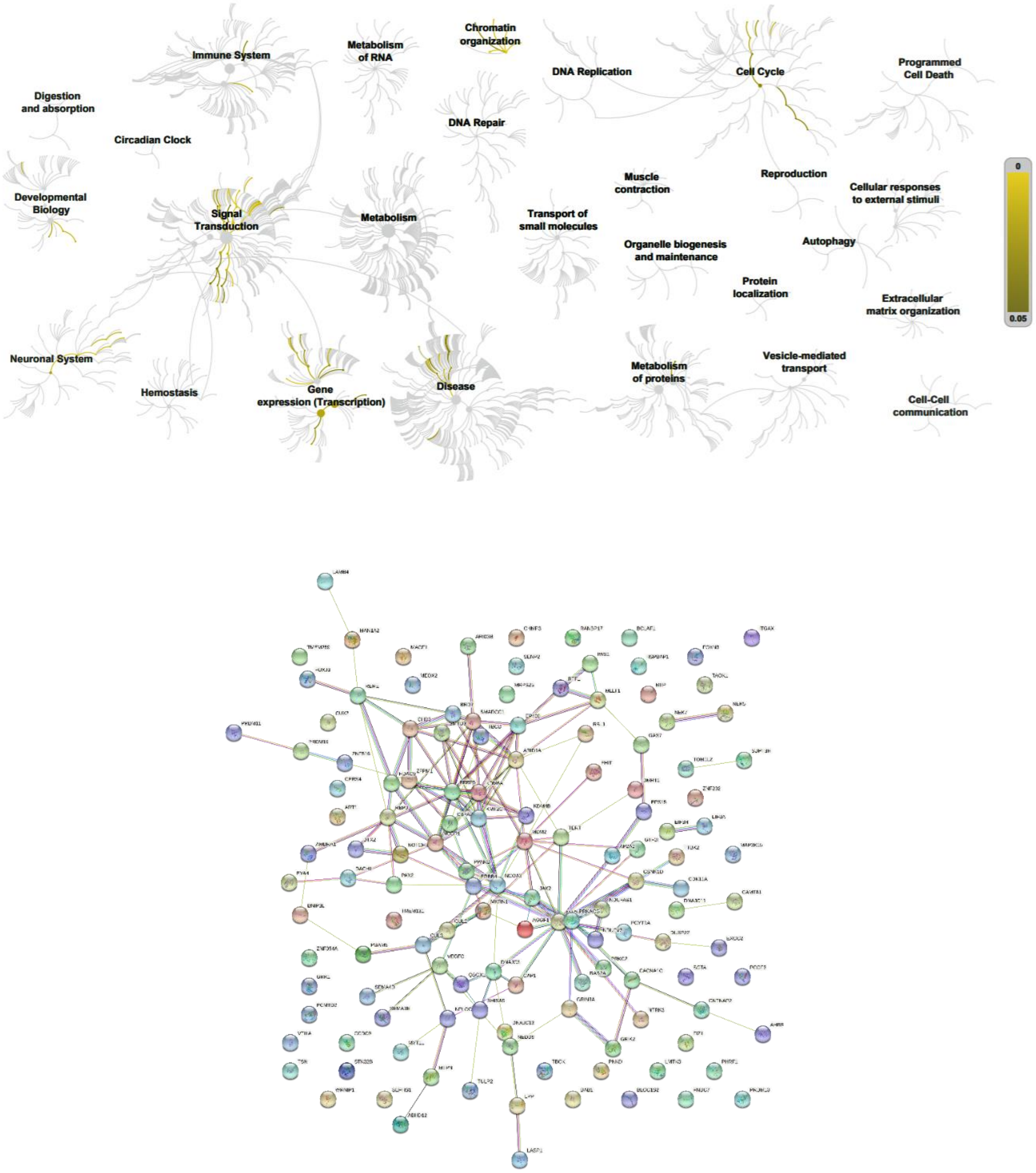


**Supplementary Figure 7.** Biochemical targets (Reactome) and pathway association (STRING) of ZPMetG-Hv2a-1B (recent) variants occurring at the CpG window region (inexact).
